# Characterizing the effect of small-scale topographic variability on co-existing native and invasive species in a heterogeneous grassland using airborne hyperspectral remote sensing

**DOI:** 10.1101/2021.04.11.439344

**Authors:** Phuong D. Dao, Alexander Axiotis, Yuhong He

## Abstract

1. Characterizing the distribution, mechanism, and behaviour of invasive species is crucial to implementing an effective plan for the protection and management of native grassland ecosystems. Hyperspectral remote sensing has been used for mapping and monitoring invasive species at various spatial and temporal scales. However, most studies focus either on invasive tree species mapping or on the landscape-level using low-spatial resolution remote sensing imagery. These low-resolution images are not fine enough to distinguish individual invasive grasses, especially in a heterogeneous environment where invasive species are small, fragmented, and co-existing with native plants with similar color and texture.
2. To capture the small yet highly dynamic invasive plants at different stages of the growing season and under various topography and hydrological conditions, we use airborne high-resolution narrow-band hyperspectral imagery (HrHSI) to map invasive species in a heterogeneous grassland ecosystem in southern Ontario, Canada.
3. The results show that there is high spectral and textural separability between invasive species and between invasive and native plants, leading to an overall species classification accuracy of up to 89.6%. The combination of resultant species-level maps and the digital elevation model (DEM) showed that seasonality is the dominant factor that drives the distribution of invasive species at the landscape level, while small-scale topographic variations partially explain local patches of invasive species.
4. This study provides insights into the feasibility of using HrHSI in mapping invasive species in a heterogeneous ecosystem and offers the means to understand the mechanism and behaviour of invasive species for a more effective grassland management strategy.

## 1 Introduction

Grasslands are spatially diverse and heterogeneous ecosystems (Banerjee et al., 2011). These ecosystems are important as they provide suitable habitat for many species (Chapman et al., 2004; Pocius et al., 2017), conserve soil and water, recycle nutrients, feed livestock (Bengtsson et al., 2019), regulate micro-climate (Wan et al., 2002), and store about 34% of the terrestrial global carbon stock (Eze et al., 2018). Native grasslands have been adversely affected by plant invasion (Gaskin et al., 2020) in the past decades. The impacts of invasive species and their interactions with native plants are complicated by varying climatic, topographic, and hydrological conditions. Invasive species may possess special traits that allow them to outcompete native plants for light, water, nutrients, and space to flourish and successfully colonize the area (Seastedt and Pyšek, 2011) in a range of environmental conditions. Mapping and monitoring the distribution, expansion, and dynamics of grasses are essential for invasive species control and management to protect native ecosystems.

Nevertheless, it is challenging to accurately map invasive grass species. Invasive grasses are typically small and often similar to native species (Ali et al., 2016), or even hybridized with native species (Stebbins, 1985). Further, grassland growth and phenology are sensitive to small-scale variation in topography. Topography is an important proxy of resource availability, microclimate, soil moisture, and edaphic site conditions (Bohlman et al., 2008; Homeier et al., 2010). For example, soils in valleys likely have higher moisture and richer nutrients than those close to the top of ridges (Gibbons and Newbery, 2003), while soils on steep slopes have lower water and nutrients availability than those in flat areas (Balvanera et al., 2011; Comita and Engelbrecht, 2009). Hence, topography and topography-controlled hydrologic regulation, as well as microclimate, drive vegetation growth and composition (Adams et al., 2014; Guisan et al., 1999), vegetation structure (Bohlman et al., 2008; Homeier et al., 2010), and plant diversity (Homeier et al., 2010; Kessler, 2002). These factors do not solely control and determine the above vegetation properties and patterns but are usually coupled with climatic conditions over the growing season. However, to the best of our knowledge, no studies have evaluated the coupled effect of these factors on the distribution, composition, and dynamics of native and invasive grass species in a heterogeneous grassland.

Remote sensing is one of the most efficient techniques for observing and monitoring the changes in vegetation species composition and health at various spatial and temporal scales. Several studies have used airborne and satellite imagery to classify grasslands from the regional scale using MODIS data (Huang et al., 2009), the landscape scale using medium resolution Landsat data (Langley et al., 2001), the community scale using high spatial resolution Quickbird data (Hall et al., 2010), to species scale using very high spatial resolution UAV images (Lu and He, 2018). However, these studies have been limited by only using a few broad multispectral bands (Marcinkowska-Ochtyra et al., 2018) or coarse spatial resolution images (Ali et al., 2016). Hence, individual species may not be classified precisely, especially when classifying species in highly mixed areas. A study by Lu and He (2018) found that species classification in a heterogeneous grassland requires decimeter-resolution imagery.

Hyperspectral remote sensing imagery (HSI) with hundreds of contiguous narrow spectral bands can distinguish subtle differences between target features (Lu et al., 2020; Marcinkowska- Ochtyra et al., 2018). Each of these narrowbands is sensitive to different plant physiological and biochemical properties and canopy structure (Lu et al., 2019). The combined use of these bands hold the potential to distinguish subtle differences between species, including species with a high degree of similarity. The HSI data have been used alone (Asner et al., 2008a) or in combination with LiDAR point cloud (Asner et al., 2008b; Marcinkowska-Ochtyra et al., 2018) to effectively distinguish native and invasive tree or grass species. However, no studies have integrated high spatial resolution HSI and DEM data to examine the seasonal, topographical, and hydrological controls on the variation and competitive ability of native and invasive species.

The spectral response of vegetation is controlled by both leaf biophysical and biochemical properties and canopy structure, including leaf pigments (e.g., chlorophyll, carotenoids, etc.), water content, nitrogen concentration, mesophyll structure, leaf thickness, leaf orientation, leaf area index (LAI), leaf angle distributions (Myneni et al., 1989), etc. The spectral characteristics associated with these plant properties may be distinct between native and invasive species, and thus assist with the identification of these species from remote sensing data. For example, Asner et al. (2008a) found distinct spectral separability between native and invasive tree species in association with leaf and canopy plant properties. However, few studies have investigated these relationships in grasslands.

In this study, airborne HrHSI is used in combination with DEM, field and laboratory vegetation measurements, and machine learning classification algorithms for mapping and characterizing the seasonal, topographical, and hydrological variations of native and invasive species in a heterogeneous grassland site in Ontario, Canada. This study aims to answer three primary research questions: (1) how do spectral and textural characteristics differ between native and invasive species? (2) to what extent can high spatial resolution hyperspectral imagery delineate native and invasive species, and characterize their intra-seasonal variability in this heterogeneous ecosystem? And (3) how much is intra-seasonality and small-scale topographic and hydrological variability driving the distribution and performance of native and invasive species?

## 2 Materials and methods

### 2.1 Study site

The experiment was conducted at the Koffler Scientific Reserve (KSR), a biological experiment site of the University of Toronto in King City, Ontario, Canada. The area is 400 m x 750 m in size, located between 44∘01’30.1” and 44∘01’43.1” in latitude and -79∘32’46.3” and -79∘32’12.3” in longitude (Fig. 1). The site is considered a heterogeneous landscape which consists of a mix of three native species, including goldenrod (*Solidago canadensis*), common milkweed (*Asclepias syriaca*), wild grape (*Vitis riparia*), and two invasive species, including smooth brome (*Bromus inermis*), orchard (*Dactylis glomerata*) (Dao et al., 2019a; Riley, 1994). These species are relatively small in size, and their colors and textures are highly similar (Ali et al., 2016). The species composition varies greatly across the growing season. In addition, from multi-year field vegetation inventory, we observe that the invasive smooth brome is less competitive over native species in lowland and flat areas but outperforms in highland, steep slopes, and dry areas. Hence, we hypothesize that there are combined effects of seasonality, topography, and hydrological condition on the distribution, composition, and spatiotemporal variation of native and invasive species in the area.

**Fig. 1.**
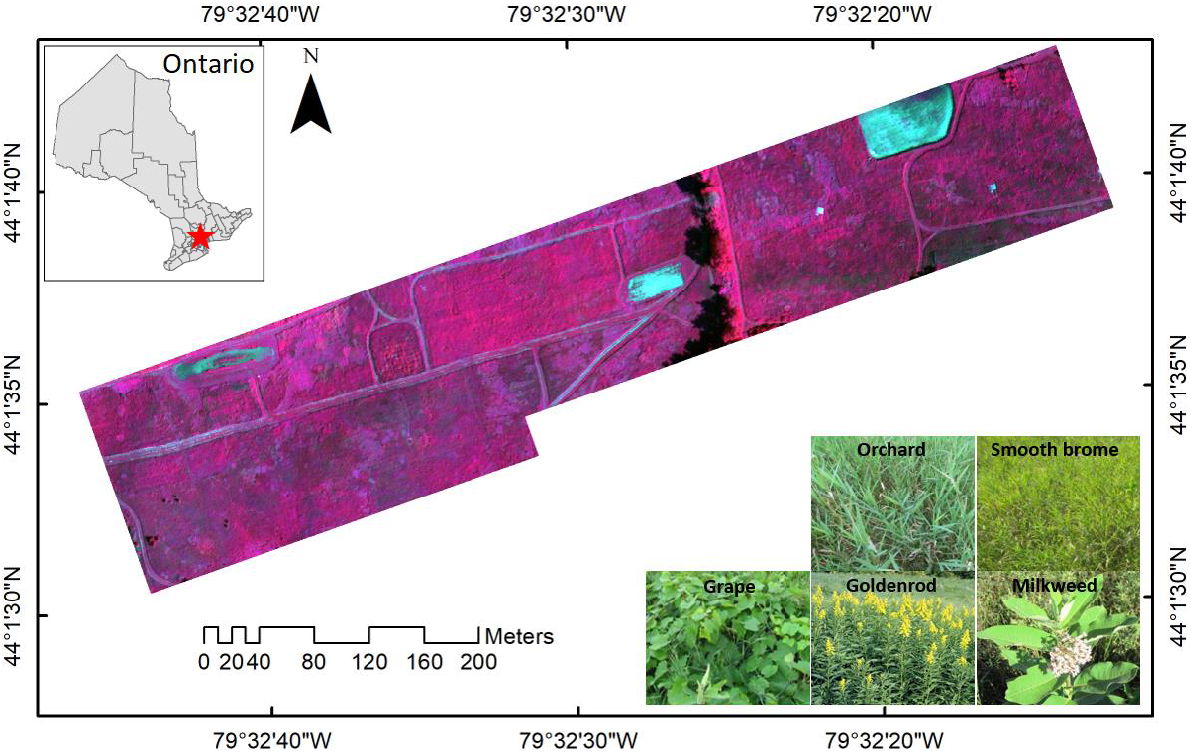
The study area with example photos of five dominant species investigated in this study.

### 2.2 Materials

#### 2.2.1 Flight mission and hyperspectral image acquisition

The Mirco-Hyperspec VNIR (A-Series) push-broom hyperspectral sensor (Fig. 2) from Headwall Photonics Inc, USA, the manned Bell 206B JetRanger helicopter, was used in this experiment. The sensor has 325 narrow spectral bands ranging from 400 nm to 1,000 nm, with a pixel depth of 12 bits. The sensor has a field of view (FOV) of 25.0403 degrees, an instantaneous field of view (IFOV) of 0.0249 degrees, and an area of interest width (the number of pixels across-track) of 1004 pixels. The sensor is equipped with an inertial measurement unit (IMU) and a global positioning system (GPS) unit that recorded the helicopter altitude and geospatial coordinates for geo-correction. The sensor comes with the HyperSpec III image acquisition software and the SpectralView image preprocessing software.

**Fig. 2.**
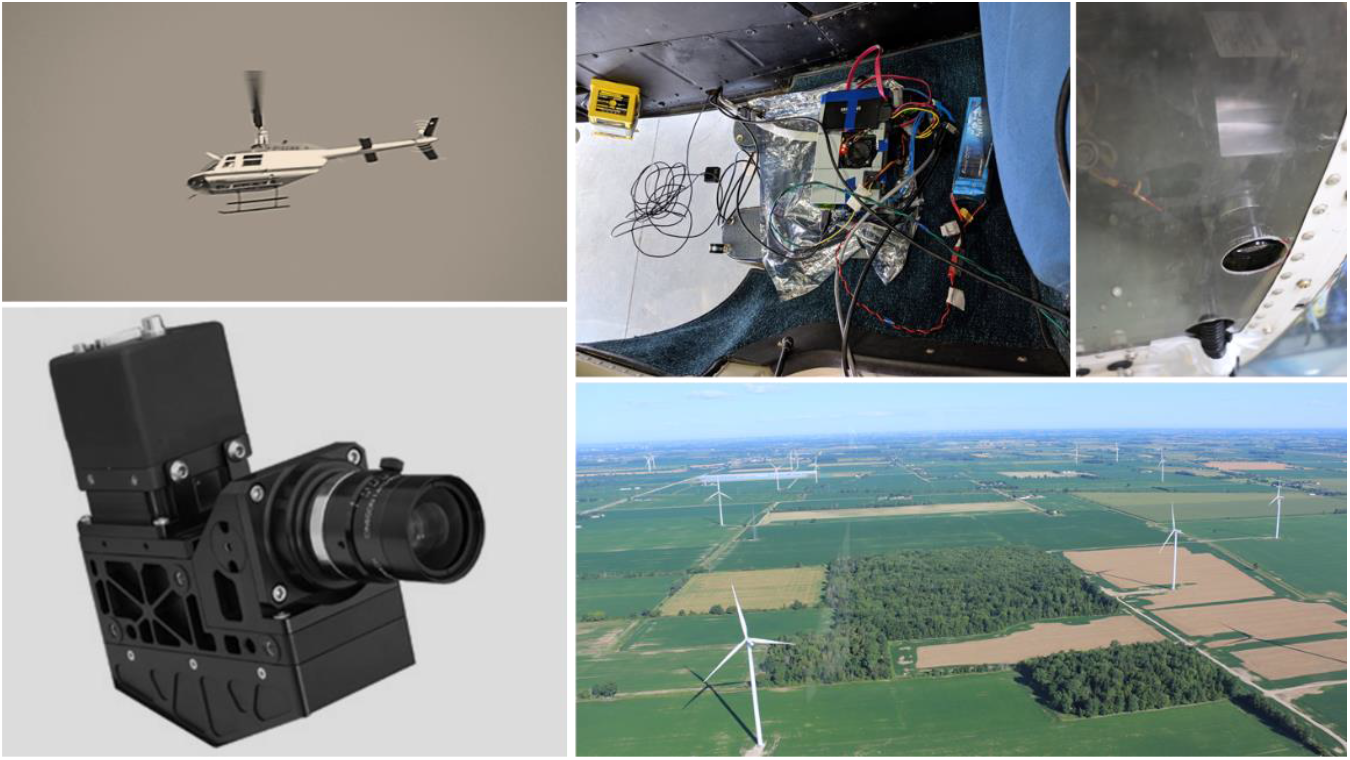
Sensor installation and flight mission. Upper left) the Bell 206B JetRanger aircraft, upper right) the micro HDPU, bottom left) the Micro-Hyperspec VNIR sensor, and bottom right a picture of the landscape from the aircraft.

In the flight mission, on 23 June 2016, 20 August 2017, and 9 September 2017, the helicopter was set to fly at an altitude about 200 m above the ground and at a speed of approximately 100 km per hour, acquiring images at a resolution of 0.20 m. The images were captured at about 10:30-11:00 am under a stable illumination. The details of the aircraft, the sensor, the acquired imagery, and the image processing software are shown in Table 1.

**Table 1.** Sensor and system specification and acquired image information.

| Item | Parameters |
| --- | --- |
| Aircraft | Bell 206B JetRanger |
| Aircraft speed | 100 km/h |
| Flight altitude | 200 m |
| Sensor | Micro-Hyperspec VNIR |
| FOV | 25.0403 degree |
| AOI width | 1004 pixels |
| Number of bands | 325 bands |
| Spectral range | 400–1000 nm |
| Spectral resolution | ~1.85 nm |
| Radiometric resolution | 12 bits |
| Spatial resolution | 0.2 m |
| Sensor software | HyperSpec III |
| Calibration software | SpectralView |

#### 2.2.2 Field reference reflectance measurement

For image radiometric calibration and atmospheric correction, ground reflectance measurement of reference targets, described in (Dao et al., 2019a), was collected at the same time of the flight mission using the FieldSpec 3 spectroradiometers (Malvern PANalytical Company, United Kingdom). The sensor provides 301 bands ranging from 400 nm to 1000 nm with an interval of 2 nm, which are the same as those of hyperspectral images and thus appropriate for calibration.

#### 2.2.3 Ground control point collection

Ground reference polygons and points were collected for image classification and accuracy assessment. The points and polygons of individual species were recorded using the real-time kinematic (RTK) technique through single-frequency (L1) EMLID RS+ GPS receivers. The signal postprocessing produced a geolocation accuracy of 0.15 m to 0.25 m. For small grass patches (with a diameter of about 2 m), the coordinates of the center points were collected, while for large homogeneous sites, polygons were recorded. In total, 372, 350, and 364 ground control points and polygons were collected for the June, August, and September images, respectively.

#### 2.2.4 Biophysical and biochemical measurements

In this study, we also examined the spectral separability in association with vegetation properties (i.e. leaf area index and chlorophyll and carotenoid contents) to assist with the species classification. Leaf area index was measured, using an AccuPAR Ceptometer (Decagon Devices, Inc.) from 25 field plots (2×2 m), at the same time as the airborne imaging campaign. At each plot, five LAI measurements were made, including ones at 4 corners and one at the centre. Three leaves for each species at each plot of the five native and invasive species were collected and stored in a cold box for laboratory measurements. A portion of each leaf sample was dried in an oven under 80oC for 24 hours to measure leaf water content per unit area. An additional portion (0.7694 cm2) of each sample was put into glassware with 4 ml of Dimethylformamide solvent and kept in a freezer under -4oC condition for 2 days before measuring the absorbance. The absorbance was measured using the GENESYS 10S UV-Vis spectrophotometer (Thermo Fisher Scientific Inc., Wisconsin, USA). The calculation of chlorophyll and carotenoid contents are expressed in Wellburn (1994) and Porra et al. (1989). LAI, together with canopy pigment and water contents, were used to link with the corresponding canopy-level spectral response.

### 2.3 Methods

The overall data interpretation process is illustrated in Fig. 3. The raw images were first preprocessed to correct the atmospheric and topographic effects, and then co-registered, mosaicked, and projected to the ground coordinates. Images were then segmented to generate image objects. Next, the spectral and textural separability of species were evaluated in association with plant leaf biochemical and canopy structural properties. Classification models were then trained and tuned with different hyperspectral and hyperspectral augmented inputs and the best model was selected for species classification. Finally, the best classification results of the three images were combined with topographic position index (TPI) and topographic wetness index (TWI) maps, calculated from a DEM, to examine the seasonal, topographic, and hydrological combined effects on vegetation dynamics and invasion.

**Fig. 3.**
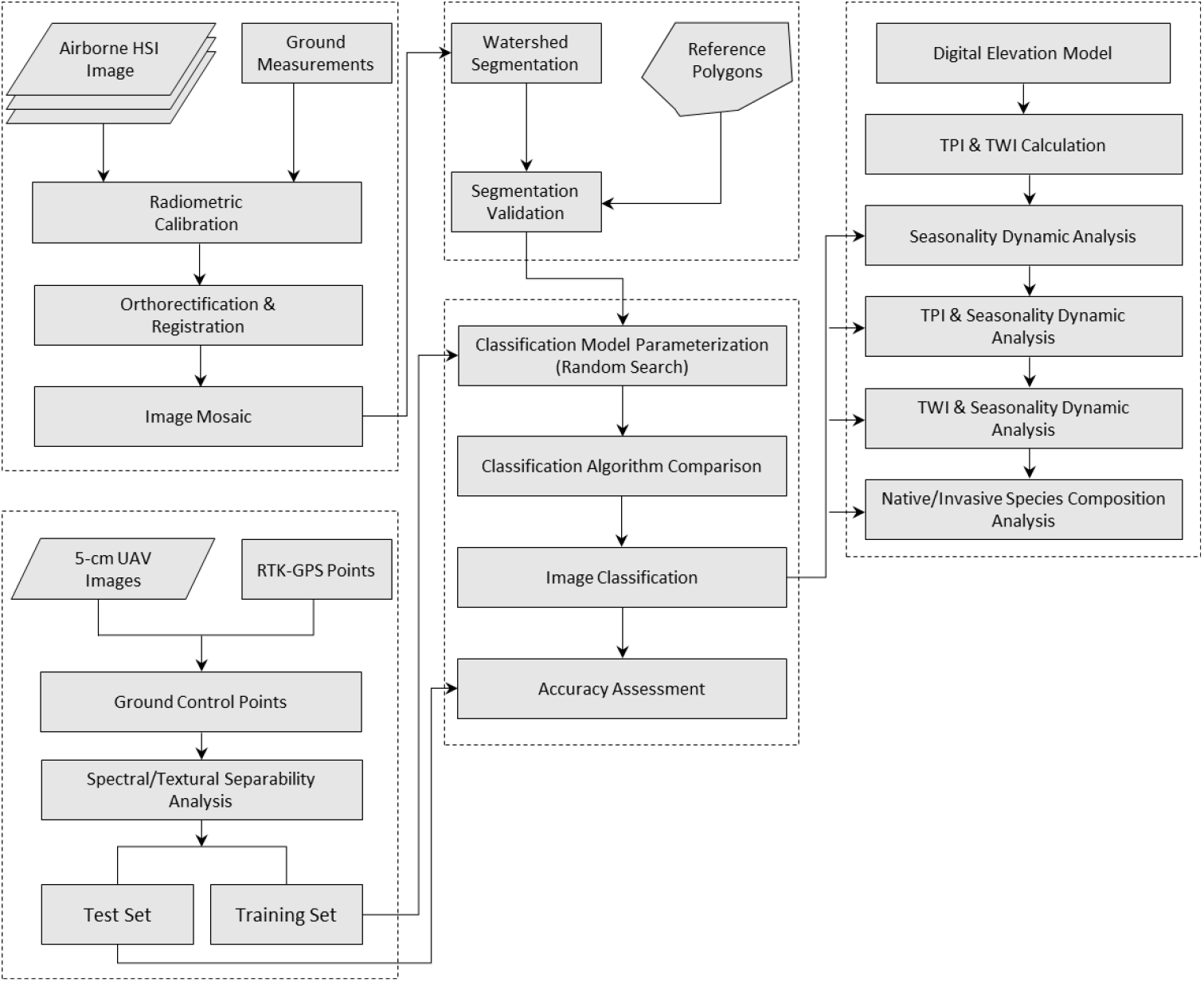
The overall design and step-by-step processes of this study.

#### 2.3.1 Image preprocessing and calibration

To reduce the geometrical distortion, the images were corrected with a digital elevation model in the SpectralView software. The collected images were projected to the North American Datum 1983 (NAD83) Universal Transverse Mercator (UTM) coordinate system. The original images of 325 narrow spectral bands were resampled to 301 bands with an interval of 2 nm. The images were co-registered using the RTK-GPS-recorded ground control points. The images were then atmospherically corrected using the calibration approach proposed by Dao et al. (2019a), to obtain surface reflectance. Samples of spectral curves of ground features, including five grass species are shown in Fig. 4.

**Fig. 4.**
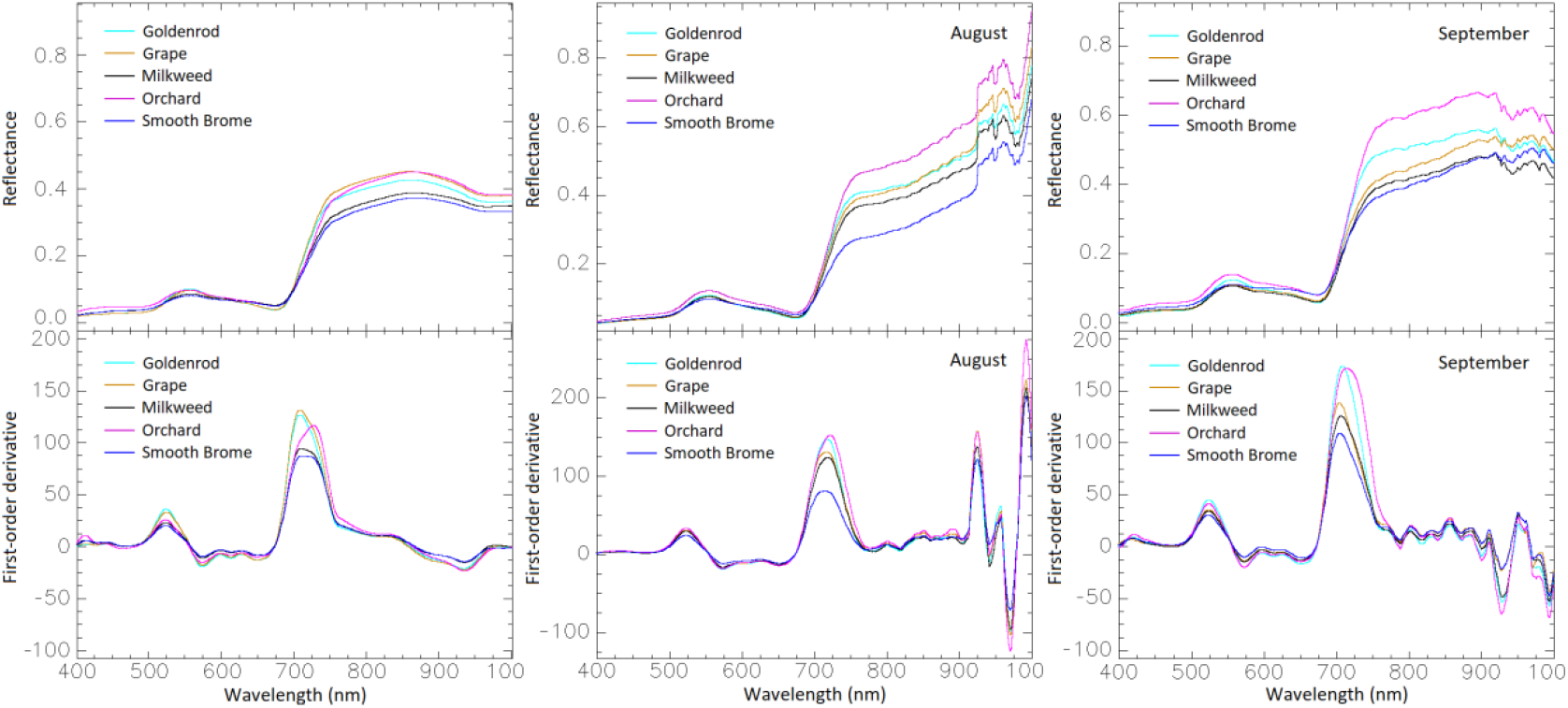
The spectral response and the first-order spectral derivative curves of five dominant species.

#### 2.3.2 Image segmentation

Image segmentation is a key step in object-based image analysis (OBIA), and has been widely used in image classification (Dao et al., 2021), owing to its ability to capture meaningful image objects with richer spatial and spectral information and minimal speckle noise (Blaschke, 2010; Dao et al., 2019b). Compared to traditional pixel-based classification methods, OBIA has proven to enhance final classification results (Dao and Liou, 2015; Dao et al., 2015). In this study, a multi-band compact watershed segmentation method (Neubert and Protzel, 2014) was used for image segmentation. The major parameters of this algorithm include the number of markers and compactness. From empirical testing and comparing with ground reference polygons, markers = 2.5×104 and compactness = 10-6 were found the best values for the three images in this study. To reduce the computation time, the segmentation was implemented on PCA-transformed images, where selected components explained 99.9% of the variance. The segments that intersected with the reference polygons (Fig. 5) were selected for the final training and test sets for classification training and validation. Specifically, segments that intersect with one or two reference polygons of the same class were selected as the training or test samples of that class, while segments that intersect with two or more polygons of different classes were ignored.

**Fig. 5.**
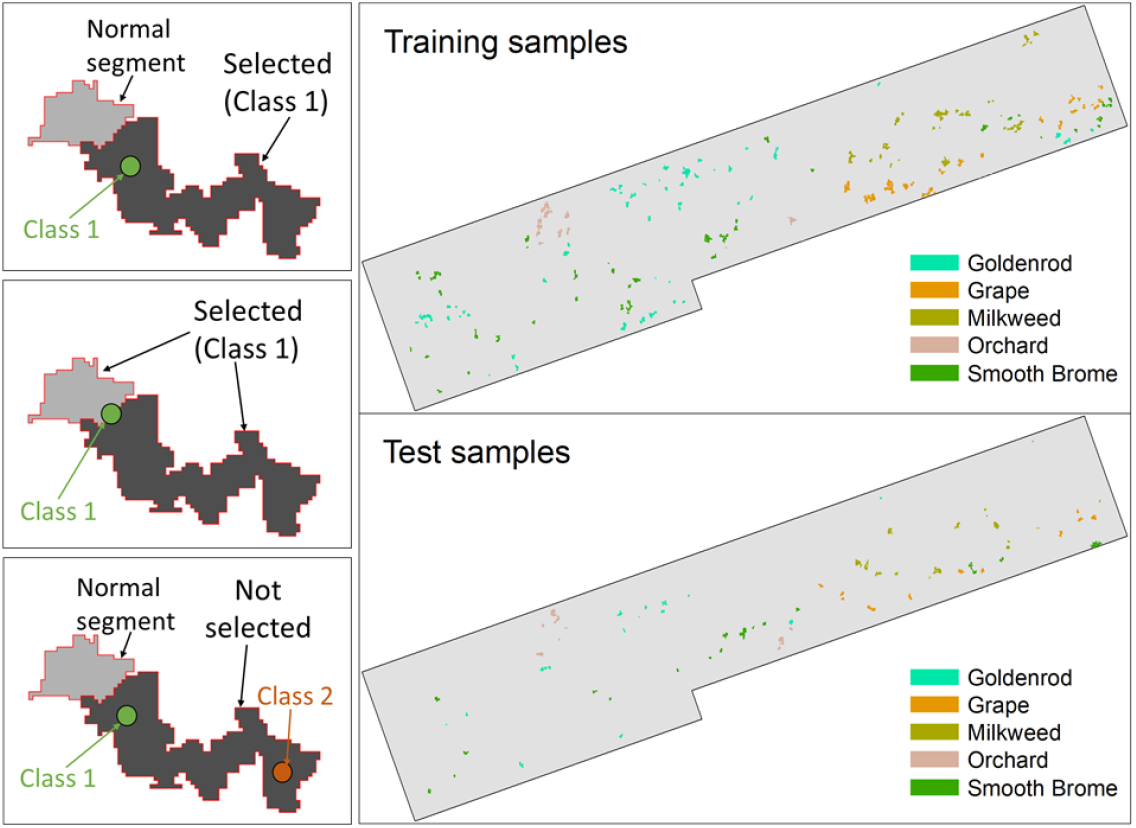
Segment sample selection procedure and training and test sample distribution.

#### 2.3.3 Species spectral separability

Species spectral separability was assessed to determine whether there are systematic spectral distinctions, between invasive species and between native and invasive species. The August image was used for demonstration as it was captured during the peak growing season. A band-by-band t-test was applied to test the differences between spectral curves, first-order spectral derivative curves, and plant properties pairs of all species. A p-value ≤ 0.05 indicates a significant difference. In addition, the Transformed Divergence (TD) (Metternicht and Zinck, 1998) spectral separability index, ranging from 0 to 2, was computed to evaluate the quality of different classification data inputs. The greater the TD value, the more spectrally separable the species are.

#### 2.3.4 Image classification

The Random Forest (RF) classification algorithm was used for classifying native and invasive species from the segmented images. RF was used for the evaluation in this study since it is a stable and reliable approach that can produce great classification results with a limited number of training samples (Ham et al., 2005; Rodriguez-Galiano et al., 2012). The major parameters in the RF algorithm in the Python 3 scikit-learn library (Pedregosa et al., 2011) are the number of trees (ntree), tree maximum depth (maxdep), number of features for the best split (mfeat), minimum samples for splitting (mspl). The selection of model inputs depends on the complexity of the scene, the spectral similarity between species. In this study, we evaluated the species classification performance of the RF algorithm with eight inputs, including full spectra (Full), full spectra and textures (Full-Tex), spectral derivatives (Deriv) (Tsai and Philpot, 2002), spectral derivative and textures (Deriv-Tex), multispectral data (Multi), multispectral data and texture (Multi-Tex), principal component analysis (PCA) transformed data (Wold et al., 1987), and PCA and textures (PCA-Tex), to determine the best model inputs. The inputs that produce the highest accuracy were used for final species classification.

For classification training, the RandomizedSearchCV tool in the Scikit-learn library in Python was used for the hyperparameter tuning and the selection of the best parameters for all the classification models. This tool randomly searches a large number of model parameter combinations and finds the best combination that maximizes the score of a k-fold cross-validation process. The combination of parameters that yields the highest validation score is then chosen for the final classification. In this paper, k = 10 was used, meaning 90% of samples were used for the searching and model fitting, and 10% of samples were used for cross-validation. We found the good ranges of parameter values for testing were: ntree = 100:2000 with a step size of 200, maxde = 10:100 with a step size of 10, nfeat = 10, 20, 100, 200, mspl = 2, 5, 10.

Reference polygons were randomly split into classification training set (80%) and the classification test set (20%). For assessment of classification performance, Producer’s Accuracy (PA), User’s Accuracy (UA), Overall Accuracy (OA) (Congalton and Green, 2008), and Cohen’s Kappa coefficient, were used (Cohen, 1960). Producer’s accuracy (0-100%) is the percentage of the reference samples in each class that is correctly classified. User’s accuracy (0-100%) is the percentage of classified image objects in each class that is actually the reference sample of the class. Overall accuracy (0-100%) is calculated by dividing the total number of correctly classified image objects by the total number of evaluated objects. Kappa coefficient (between -1 and 1) presents the agreement between the classification results and the reference data. (McHugh, 2012).

#### 2.3.5 Species analysis across TPI and TWI gradients

Topographic position index 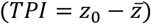, derived from a DEM, measures the difference between the elevation of the central point (*z*_0_) and the mean elevation 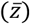 of a predefined neighbourhood–with a radius *R* (De Reu et al., 2013; Weiss, 2001). The values of TPI are between -1 and 1, which represent topographic relief from valleys, flat land to ridges. We calculated TPI using an 8-m DEM through a 21 by 21 cell window that captures localized topographic variation and maintains diverging and converging slopes in the landscape. According to Weiss (2001), TPI was classified into six categories, including ridge, upper slope, middle slope, flat area, lower slope, and valley, in Table 2, where *SD* is the standard deviation of elevation within the neighbourhood. Topographic wetness index (TWI) (Beven and Kirkby, 1979), calculated from a DEM, is a proxy of soil moisture, groundwater availability, and nutrient richness. The index is calculated as *TWI* = *ln*(*a*/tan (*β*)), where *a* is the specific catchment area and *β* is the slope of the terrain. To categorize TWI into meaningful subbasins ground measurements of soil type and soil characteristics are required. Since these measurements are not available in this study, we classified the TWI into six generic categories (Table 2), from TWI-1 to TWI-6, using the natural-break classification method. The lower the category numbers, the drier the area, while the higher the category numbers, the wetter the area.

**Table 2.** The categorization schemes of the TPI and TWI indices.

| TPI category | Criterion | TWI category | Value range |
| --- | --- | --- | --- |
| Ridge | $z_0 > 1SD$ | TWI-1 | $TWI \leq 70$ |
| Upper slope | $1SD \geq z_0 > 0.5SD$ | TWI-2 | $70 < TWI \leq 215$ |
| Middle slope | $0.5SD \geq z_0 \geq -0.5SD, \text{slope} > 5^\circ$ | TWI-3 | $215 < TWI \leq 430$ |
| Flat area | $0.5SD \geq z_0 \geq -0.5SD, \text{slope} \leq 5^\circ$ | TWI-4 | $430 < TWI \leq 890$ |
| Toe slope | $-0.5SD > z_0 \geq -1SD$ | TWI-5 | $890 < TWI \leq 1540$ |
| Valley | $z_0 < -1SD$ | TWI-6 | $1540 < TWI$ |

These indices have been widely used as means to study the effect of topography on vegetation pattern, productivity, composition, and diversity (Adams et al., 2014; Alexander et al., 2016; Guisan et al., 1999; Zinko et al., 2005). In this study, TPI and TWI maps were joined with species classification maps to investigate the small-scale combined effect of topography and seasonality on the distribution, composition, and performance of co-occurring native and invasive grass species in the area. Sample transact profiles with annotations of six TPI and TWI categories are illustrated in Fig. 6.

**Fig. 6.**
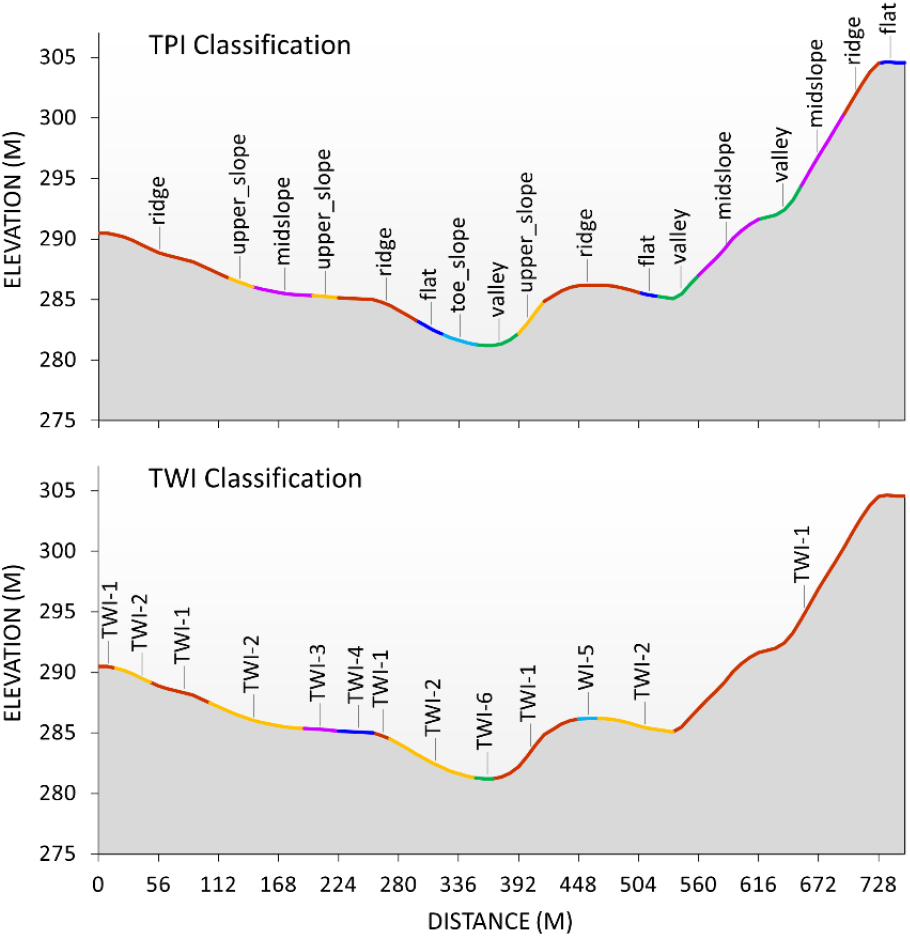
A profile graph of the digital elevation with labeled TPI and TWI segments of the study site.

## 3 Results and discussion

### 3.1 Species spectral and textural separability

Species spectral analysis in association with leaf biochemical and canopy structural properties is illustrated in Fig. 7. For the significance test of spectral response, the bands with significant differences were highlighted using grey bars. Similarly, the significance test results for the pairs of plant properties of all species are shown in the rectangle grid (in the bottom boxplots), in which the rectangles of statistical significance are highlighted in grey. The t-test shows that the spectral reflectance of the invasive smooth brome and orchard were significantly different from those of the other native species, with orchard showing the highest reflectance, especially in the blue (below 500 nm), red-edge (680-730 nm), and near-infrared (NIR, beyond 730 nm) regions. Similar patterns were also observed in the test results of the spectral derivative, except in the long-wavelength NIR region (800-1000 nm). These spectral separability results are well aligned with the significance tests of the chlorophyll, carotenoid, water content, and leaf area index, in which at least two out of four properties of smooth brome and orchard were statistically different from those of other native species. This indicated that systematic spectral differences exist between native and invasive species in the area.

**Fig. 7.**
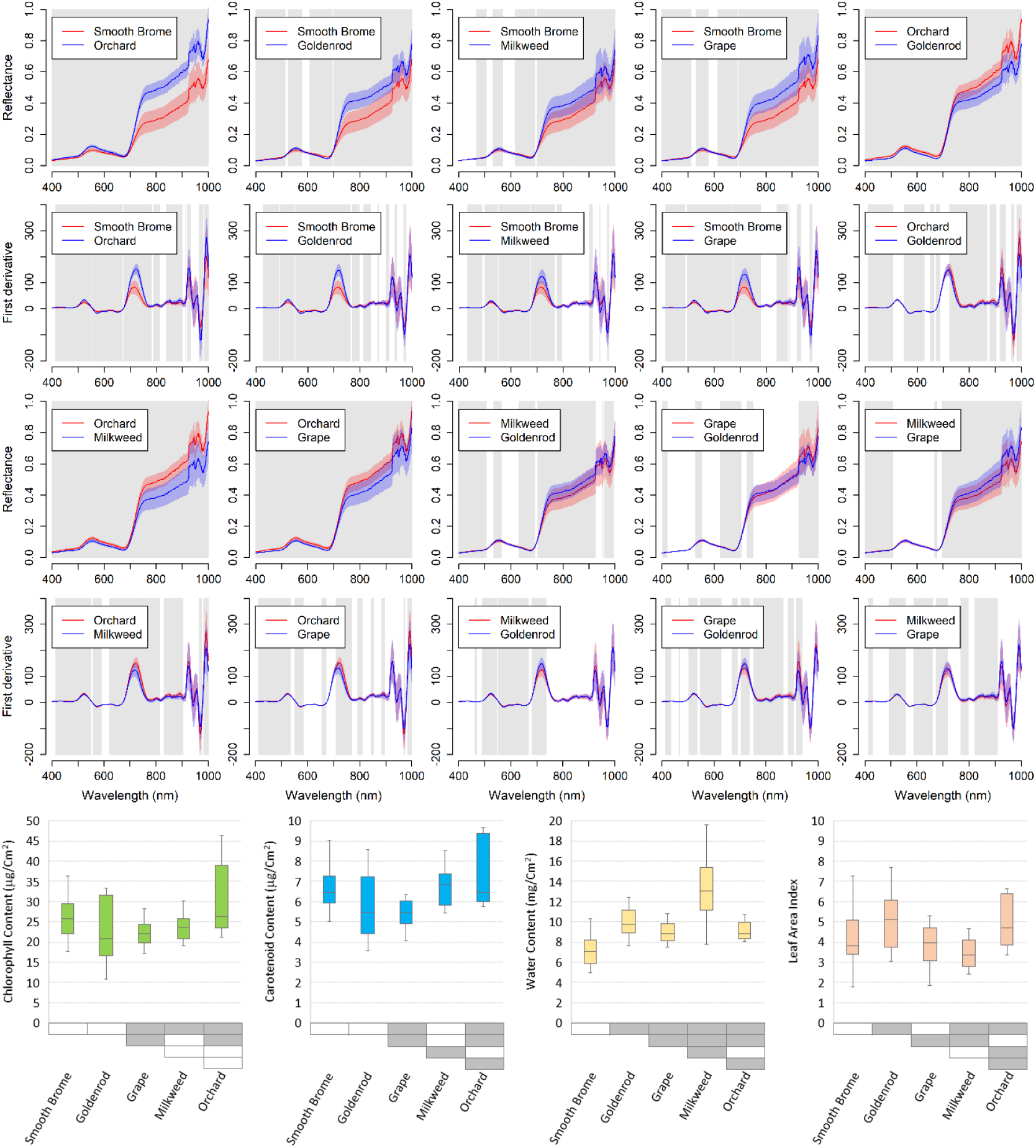
The spectral separability between all pairs of native and invasive species in association with plant properties. The grey color in the spectral response figures indicates a significant difference. The rectangle grids at the bottom boxplots show the plant property test results; grey rectangles are the pairs of significant differences (p-value ≤ 0.05).

The t-test showed that three native counterparts (goldenrod, milkweed, and grape) had a considerably high spectral similarity, especially the green and red regions (500-680 nm). Consistently, no significant differences were observed between native species’ chlorophyll content, which controls the light absorption of leaves in these regions. This spectral similarity poses challenges to classify these native species in remote sensing images.

It is important to note that the t-test results showed that the spectral responses of a pair of species might be statistically different in one spectral region. In contrast, those of other pairs were different in the other spectral regions. This suggests that if a researcher only used a few spectral bands to map these species, it could lead to misclassification between species, especially between native species.

The spectral separability measures of different inputs from the three images are shown in Table 3, and example images of textures are shown in Fig. A1. The use of full spectra and spectral derivative effectively delineated five species (TD = 1.59-2.00 with full spectra and TD = 1.41-2.00 with spectral derivative). We expected the use of these inputs to produce the most accurate and reliable species classification results. In contrast, the spectral separability decreased substantially when using multispectra (TD = 0.35-1.77), PCA (TD = 0.09-1.53), or texture (TD = 0.01-2.00). The use of these data may result in misclassification between spectrally similar species. Some textural information, including variance, homogeneity, contrast, dissimilarity, entropy, and correlation (Farwell et al., 2020), are distinct among some species between growth stages, and the inclusion of this information may improve the identification of these species.

**Table 3.** Species spectral and textural separability (TD value) with different inputs from the three images calculated from training samples.

|  | June Image |  |  |  | August Image |  |  |  | September Image |  |  |  |
| --- | --- | --- | --- | --- | --- | --- | --- | --- | --- | --- | --- | --- |
|  | Goldenrod | Orchard | Milkweed | Grape | Goldenrod | Orchard | Milkweed | Grape | Goldenrod | Orchard | Milkweed | Grape |
| **Full spectra** |  |  |  |  |  |  |  |  |  |  |  |  |
| Smooth brome | 1.86 | 1.92 | 1.96 | 1.99 | 1.83 | 1.89 | 1.59 | 1.83 | 1.96 | 1.99 | 1.87 | 1.96 |
| Goldenrod |  | 1.97 | 2.00 | 2.00 |  | 1.85 | 1.71 | 1.81 |  | 2.00 | 1.95 | 1.99 |
| Orchard |  |  | 2.00 | 2.00 |  |  | 1.88 | 1.97 |  |  | 2.00 | 2.00 |
| Milkweed |  |  |  | 1.79 |  |  |  | 1.63 |  |  |  | 1.95 |
| **Spectral derivative** |  |  |  |  |  |  |  |  |  |  |  |  |
| Smooth brome | 1.83 | 1.86 | 1.92 | 1.96 | 1.75 | 1.84 | 1.44 | 1.69 | 1.88 | 1.99 | 1.69 | 1.87 |
| Goldenrod |  | 1.95 | 2.00 | 2.00 |  | 1.74 | 1.53 | 1.74 |  | 1.99 | 1.77 | 1.96 |
| Orchard |  |  | 2.00 | 2.00 |  |  | 1.76 | 1.89 |  |  | 1.98 | 2.00 |
| Milkweed |  |  |  | 1.75 |  |  |  | 1.41 |  |  |  | 1.82 |
| **Multispectral** |  |  |  |  |  |  |  |  |  |  |  |  |
| Smooth brome | 1.66 | 0.72 | 0.35 | 1.59 | 1.18 | 0.98 | 0.48 | 1.09 | 1.24 | 0.59 | 0.44 | 0.90 |
| Goldenrod |  | 1.32 | 1.77 | 0.73 |  | 0.79 | 0.79 | 0.44 |  | 1.47 | 0.90 | 0.57 |
| Orchard |  |  | 0.79 | 1.68 |  |  | 1.01 | 1.28 |  |  | 1.04 | 1.29 |
| Milkweed |  |  |  | 1.29 |  |  |  | 0.39 |  |  |  | 0.53 |
| **PCA** |  |  |  |  |  |  |  |  |  |  |  |  |
| Smooth brome | 1.45 | 1.16 | 0.42 | 1.53 | 0.94 | 0.77 | 0.53 | 0.75 | 1.34 | 1.43 | 0.94 | 0.95 |
| Goldenrod |  | 1.34 | 0.87 | 0.32 |  | 0.36 | 0.41 | 0.31 |  | 1.15 | 0.44 | 0.75 |
| Orchard |  |  | 1.04 | 1.38 |  |  | 0.74 | 0.60 |  |  | 1.28 | 0.69 |
| Milkweed |  |  |  | 0.89 |  |  |  | 0.09 |  |  |  | 0.52 |
| **Texture** |  |  |  |  |  |  |  |  |  |  |  |  |
| Smooth brome | 1.94 | 2.00 | 1.59 | 2.00 | 0.67 | 2.00 | 0.61 | 2.00 | 2.00 | 2.00 | 0.53 | 2.00 |
| Goldenrod |  | 2.00 | 0.32 | 1.08 |  | 1.99 | 0.01 | 2.00 |  | 2.00 | 2.00 | 0.58 |
| Orchard |  |  | 2.00 | 2.00 |  |  | 1.99 | 0.19 |  |  | 1.91 | 2.00 |
| Milkweed |  |  |  | 1.21 |  |  |  | 2.00 |  |  |  | 2.00 |

### 3.2 Species classification results

The comparison of the RF classifier of the eight inputs for the three images is shown in Table 4. Models with Deriv and Full inputs always produced the highest accuracy, especially when combined with textures (OA = 84.1-89.5% and Kapp = 0.796-0.865 for June image, OA = 83.9-89.6% and Kappa = 0.797-0.870 for August image, and OA = 80.9-83.0% and Kapp = 0.756-0.785 for September image). The accuracy of the Multi, Multi-Tex, PCA, PCA-Tex models were lower. These results well-reflect the spectral separability between species in Table 3. The inclusion of image textures improved the classification accuracy of the August and September images, but not that of the June image. The September classification accuracy was the lowest, possibly due to the mixture of senesced and dried vegetation and the background litter. In terms of computation, the Full, Full-Text, Deriv, and Deriv-Tex models with greater numbers of input features were more computationally expensive (required 161-639 seconds) than the other four models (60-111 seconds). However, the high computation times are not alarming in hyperspectral data analysis, considering their significant improvement in classification accuracy. The data inputs (Deriv for the June image and Deriv-Tex for the August and September images), with the highest accuracy, were used for the final species classification.

**Table 4.**
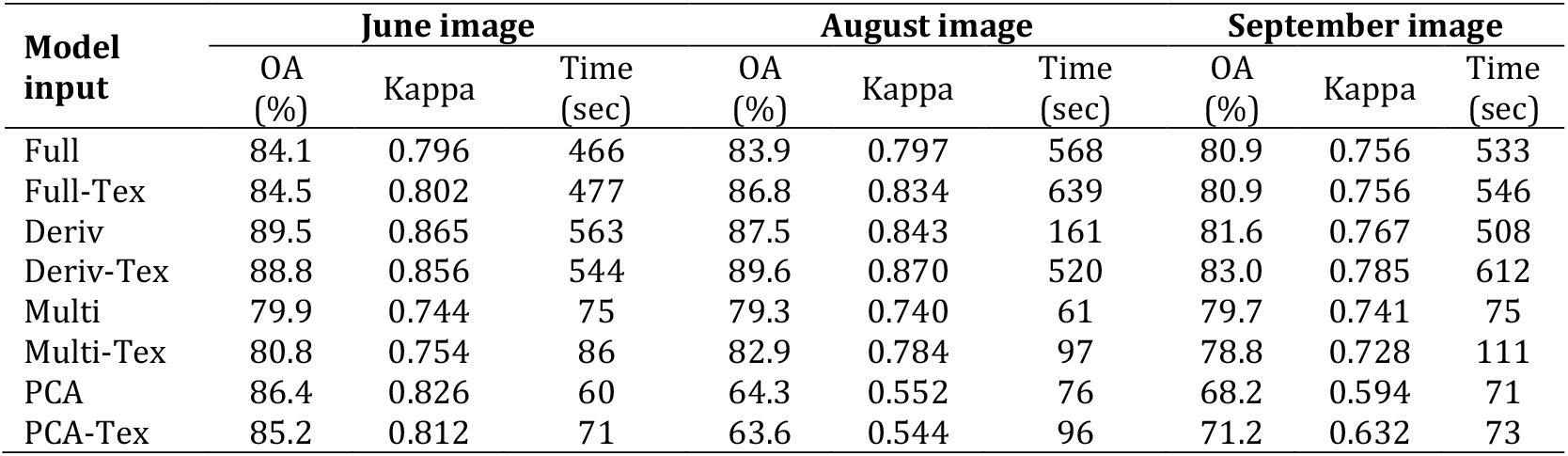
The performance and accuracy of the RF classifier with different inputs of the thee images.

The species classification accuracy (Fig. 8) of the RF algorithm was evaluated with different model inputs (Full, Full-Tex, Deriv, Deriv-Tex, Multi, Multi-Tex, PCA, and PCA-Tex). RF performed the best when Deriv-Tex input was used, followed by Full, Multi, and PCA inputs. When textural information was combined with the above inputs, the accuracy was improved for the August and September images, especially for smooth brome and milkweed. However, it did not enhance the results of the June image. These results suggest that, when limited spectral information (Multi or PCA) was used, the classification accuracy of some species could be improved by adding textural information.

**Fig. 8.**
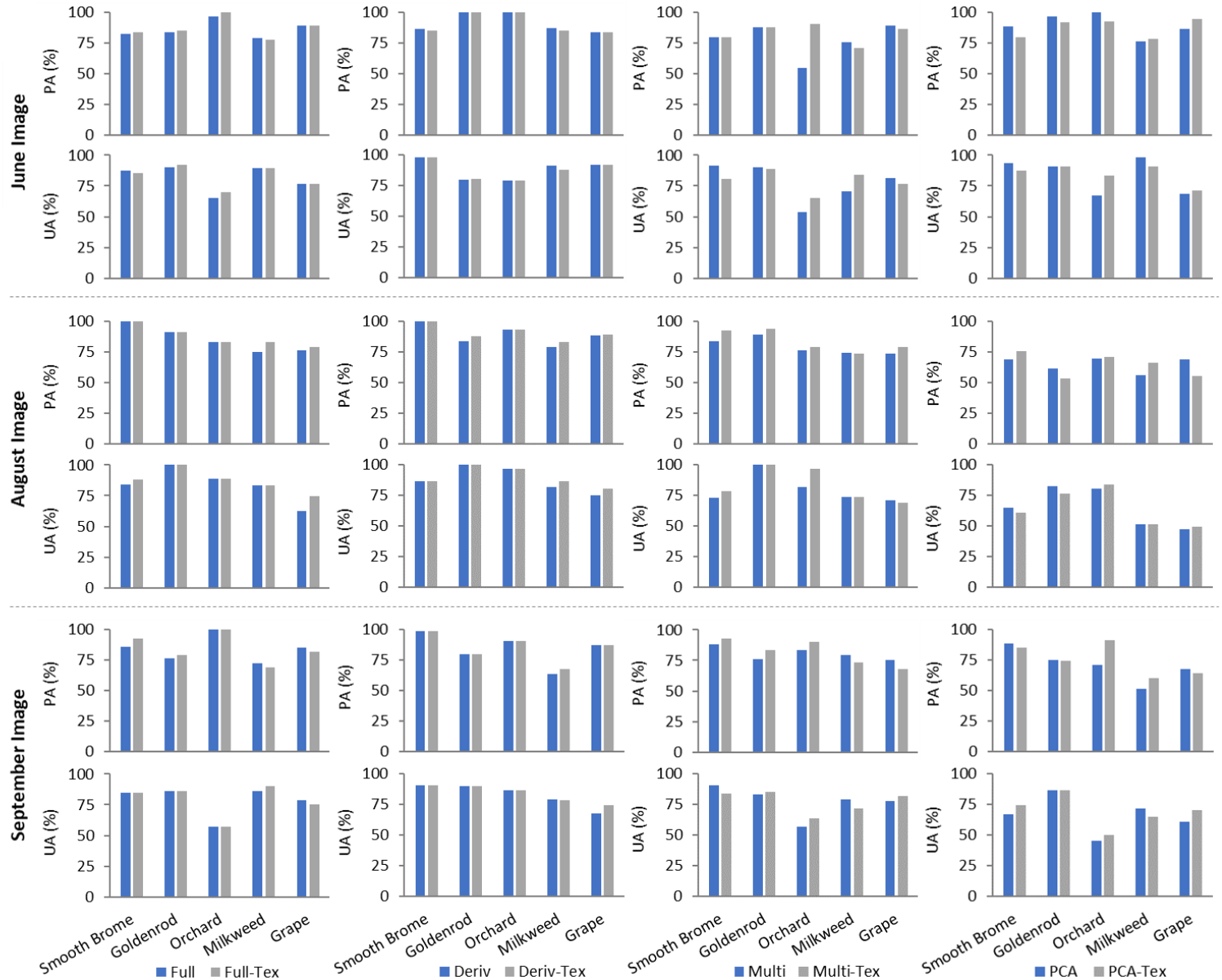
Comparing the classification of species when textures are included and not included. From top to bottom are the figures of producer’s accuracy (left figures) and user’s accuracy of Full and Full-Tex, Deriv and Deriv-Tex, Multi and Multi-Tex, and PCA and PCA-Tex models, respectively.

The detailed RF classification accuracies of the three images, when the optimal data inputs used, are shown in Table 5. Overall, the classification achieved high accuracy in all three images, with the highest for August image (OA = 89.6%, Kappa = 0.870), followed by June image (OA = 89.5%, Kappa = 0.865) and the September image (OA = 83.0%, Kappa = 0.785). In September, PAs and Uas decreased in goldenrod (PA from above 87.9% August to 79.6% in September and UA from 100% in August to 89.7% in September), milkweed (PA from 82.7% in August to 67.3% in September and UA from 86.1% in August to 78.3% in September), and in grape (PA from 89.1% in August to 87.0% in September and UA from 80.4% in August to 74.1% in September). This is expected since senesced leaves in the late growing season were more difficult to capture, and sparsely vegetated areas could be misclassified to background soil and litter.

**Table 5.** The accuracy assessment of Random Forest classification of three images with the best inputs (Deriv for June image, and Deriv-Tex for August and September images).

| Reference | Predicted |  |  |  |  |  |
| --- | --- | --- | --- | --- | --- | --- |
|  | Smooth brome | Goldenrod | Orchard | Milkweed | Grape | UA |
| **23 June 2016** (OA = 89.5%, Kappa = 0.865) |  |  |  |  |  |  |
| Smooth brome | 95 | 0 | 0 | 2 | 0 | 97.9% |
| Goldenrod | 0 | 71 | 0 | 4 | 14 | 79.8% |
| Orchard | 2 | 0 | 34 | 7 | 0 | 79.1% |
| Milkweed | 7 | 0 | 0 | 99 | 2 | 91.7% |
| Grape | 6 | 0 | 0 | 1 | 83 | 92.2% |
| PA | 86.4% | 100% | 100% | 87.6% | 83.8% |  |
| **20 August 2017** (OA = 89.6%, Kappa = 0.870) |  |  |  |  |  |  |
| Smooth brome | 44 | 0 | 0 | 7 | 0 | 86.3% |
| Goldenrod | 0 | 51 | 0 | 0 | 0 | 100% |
| Orchard | 0 | 0 | 53 | 0 | 2 | 96.4% |
| Milkweed | 0 | 7 | 0 | 62 | 3 | 86.1% |
| Grape | 0 | 0 | 4 | 6 | 41 | 80.4% |
| PA | 100% | 87.9% | 93.0% | 82.7% | 89.1% |  |
| **9 September 2017** (OA = 83.0%, Kappa = 0.785) |  |  |  |  |  |  |
| Smooth brome | 84 | 1 | 4 | 4 | 0 | 90.3% |
| Goldenrod | 1 | 78 | 0 | 8 | 0 | 89.7% |
| Orchard | 0 | 0 | 38 | 2 | 4 | 86.4% |
| Milkweed | 0 | 12 | 0 | 72 | 8 | 78.3% |
| Grape | 0 | 7 | 0 | 21 | 80 | 74.1% |
| PA | 98.8% | 79.6% | 90.5% | 67.3% | 87.0% |  |

To confirm that hyperspectral imagery outperforms multispectral imagery in species classification, the results (of August image) were compared to that of a previous study (Lu and He, 2018), which used UAV-acquired multispectral imagery in the same study area with identical spatial resolution (0.20 m) and the same classification algorithm–Random Forest. Our overall accuracy with hyperspectral data was 89.6%, whereas the previous study had achieved 73% using three-band data.

### 3.3 Seasonal species dynamic analysis

The final species classification maps with optimal inputs of three images are shown in Fig. 9, and the proportions of species across the entire study area are shown in Fig. 10. In June, invasive smooth brome dominated almost all areas with various elevations and slopes (57% of total area). The dominant distribution of smooth brome is expected because ground survey indicated that the species is a cool-season grass that often grows early in the season, which was also discussed by Stubbendieck et al. (1992). Orchard, the other invasive species, occupied only a few small lowland regions (5%). It is expected because orchard is more susceptible to competition (Muyt, 2001). The second most dominant species was the native goldenrod (21%) that also grows earlier in the growing season (Weber and Schmid, 1998); the species mainly occupied the lowland and flat areas in the centre of the site. Other plants that were seen in the field site were milkweed (11%) and grape (6%); however, these native plants only grew sparsely in lowland, flat, and wet areas.

**Fig. 9.**
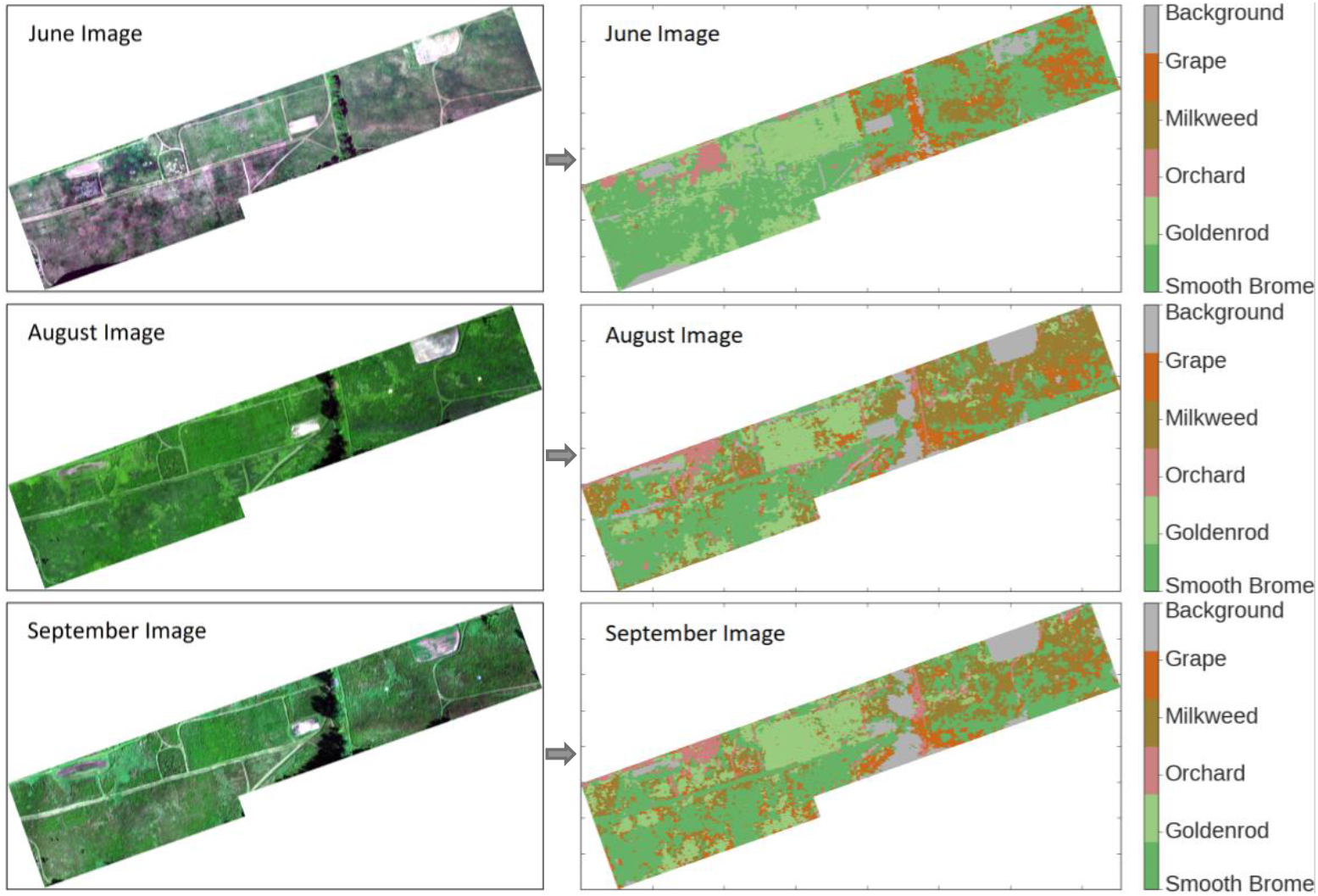
Classification results of the June, August, and September images without textures.

**Fig. 10.**
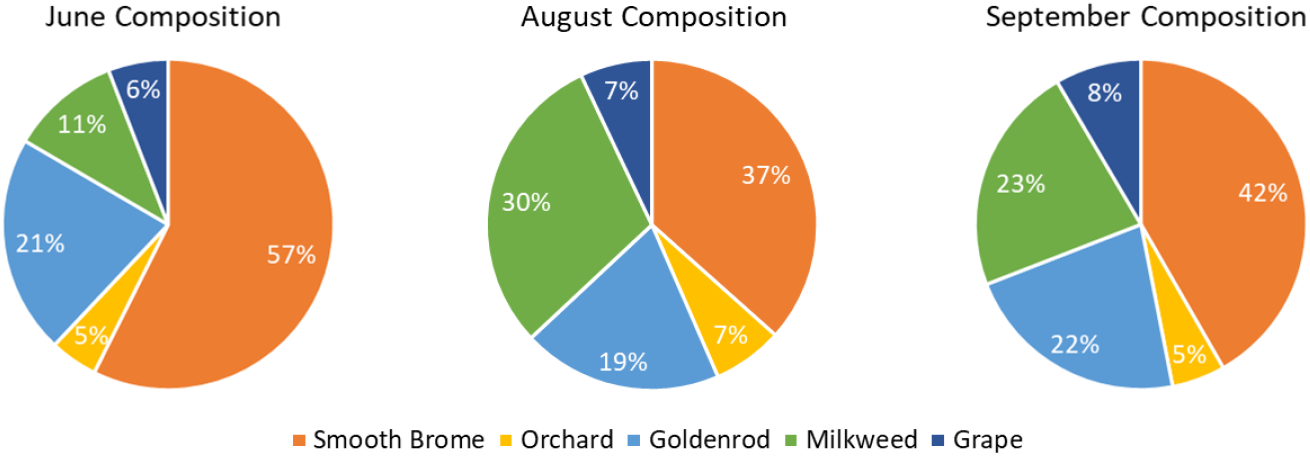
Percentages of species and in the entire study site in June, August, and September.

In August, both the species composition and distribution changed substantially. Smooth brome was still the most dominant (37% of total area), followed by milkweed (30%) and goldenrod (19%). Orchard and grape seemed to remain stable. Compared to June, goldenrod and milkweed grew dominantly and even outcompeted invasive smooth brome in lowland and flat regions. However, smooth brome Outcompeted native species in high elevation, steep slope, and dry areas. This is expected as smooth brome is a drought-tolerant species (Sheaffer et al., 1992). Orchard was the least dominant species in August as it is less aggressive and the least persistent in dry environments (Sheaffer et al., 1992).

In September, the area of grape and milkweed (both were in their late growing season) decreased to a moderate extent while the area of smooth brome increased substantially from 37% to 42%. Interestingly, the area of smooth brome increased in lowland regions that were previously occupied by milkweed and grape but now experiencing senescence. This is expected as smooth brome either stayed green for longer or experienced fall green-up (Stubbendieck et al., 1992) when the environmental conditions are favorable, such as when there is sufficient soil moisture. The area of the goldenrod and orchard did not change much as their life spans are longer and still grew well in the mid of September (Weber and Schmid, 1998).

### 3.4 Species dynamics analysis across TPI and TWI gradients

The species composition over the topographic regions and growing seasons is shown in Fig. 11. The invasive smooth brome grew and dominated all the topographic regions (higher than 50% in five out of six areas) in June, with the greatest proportion in ridges (approximately 74%). Compared to its area across the study site, the highest proportion of smooth brome was also found in ridges (over 39%). This is reasonable as smooth brome is a cool-season C3 grass that usually grows earlier in the season (Stubbendieck et al., 1992).

**Fig. 11.**
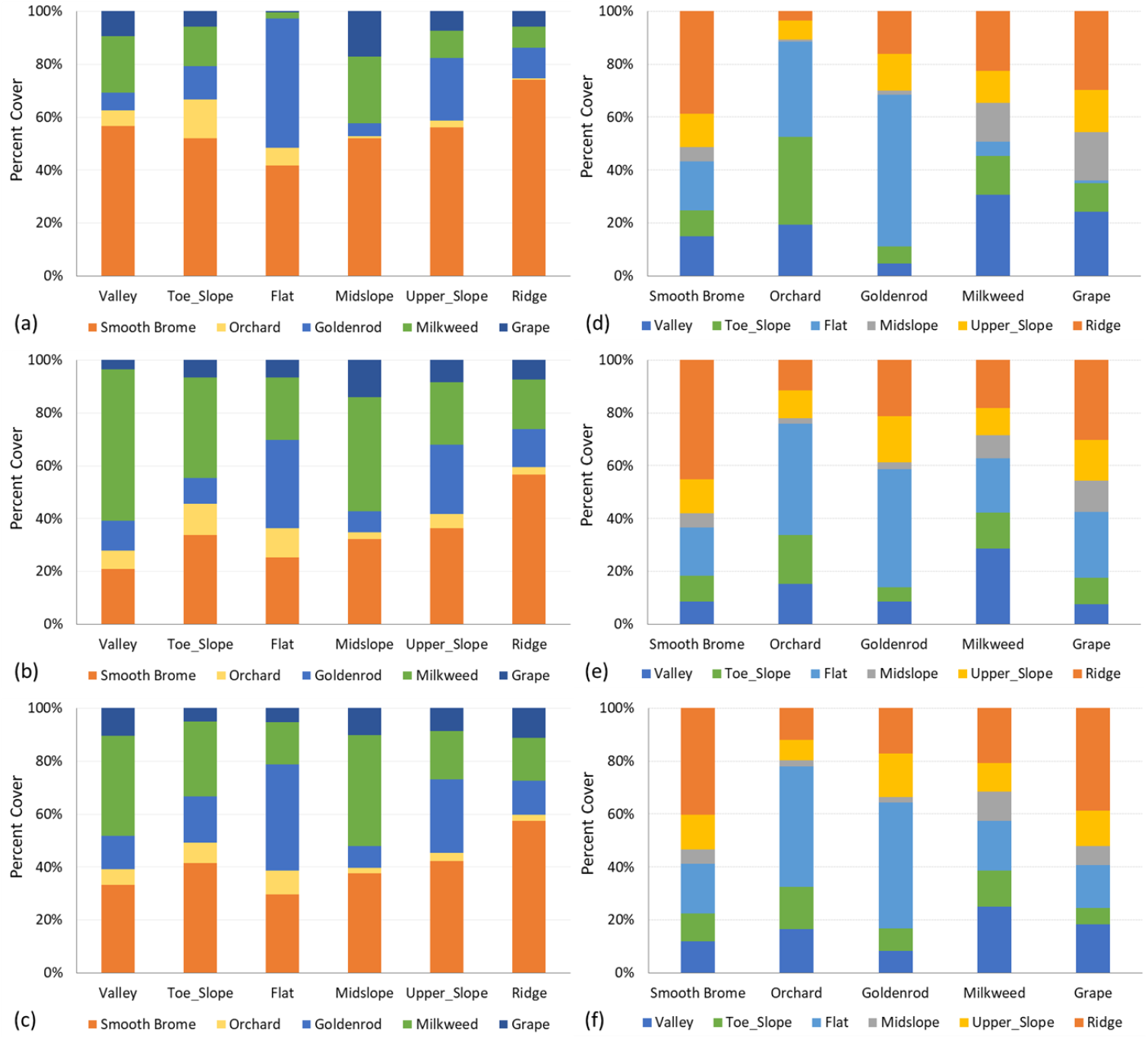
Species percentages in different TPI categories. (a), (b), (c) are species percent cover over the total area of all species in each topographic region in June, August, and September, respectively. (d), (e), (f) are species percent cover in each topographic region over its total site area in June, August, and September, respectively.

In August, however, the percentage cover of smooth brome decreased substantially, especially in lowland and flat regions. In the valleys, toe slope, and flat regions, it decreased from 57%, 52%, 42% in June to 21%, 33%, 25% in August, respectively. However, smooth brome remained the most dominant species in upper slopes (36%) and ridges (57%). This is especially evident on ridges where 45% of smooth brome was found. In contrast, the areas of milkweed increased considerably in all topographic regions, especially in valley and toe slope, in August. Whilst, grape, orchard, and goldenrod experienced minimal changes between the two months.

In September, the percentage cover of the invasive smooth brome increased by 2-12% in all topographic regions, and that of goldenrod increased by 1-8% in five out of six regions. In contrast, the area of milkweed decreased (by 2-19%) significantly in all topographic regions, especially in valleys, with a 19% decrease. The increase of the earlier two species and the reduction of the latter are explainable. First, smooth brome that occupied the lowland and flat areas in June were outgrown and co-existed, but not completely replaced by milkweed in August. However, in September, smooth brome persisted in the area while milkweed was at its senescing stage. Second, smooth brome experienced a fall green-up, mainly in lowland and flat areas with favourable conditions, especially due to high precipitation in 2017. In general, despite variation across the growing seasons, smooth brome and grape mostly appeared in ridges, while orchard, goldenrod, and milkweed were primarily seen in flat and upper slope regions.

The species composition over six topographic wetness categories is demonstrated in Fig 12. It is evident that in June, the invasive smooth brome was the most dominant species in the three driest regions TWI-1, TWI-2, and TWI-3, with the percentage covers of 61%, 62%, and 44%, respectively, indicating its intention to reside and dominate areas with dry soil. In contrast, goldenrod dominantly occupied the three wettest regions TWI-4, TWI-5, and TWI-6, with the percent covers of 51%, 59%, and 90%, respectively. Across all TWI categories, smooth brome, grape, and milkweed had their largest proportion in the TWI-1 class, while orchard and goldenrod had their largest proportion in the TWI-2 category. In August, despite the decrease in the percent cover, smooth brome remained the most dominant species in dry regions while goldenrod was dominant in wetter regions. Milkweed expanded considerably in all wetness categories. Similar to the patterns in the TPIs, the area of smooth brome increased moderately in most TWI categories, especially in the wet regions TWI-5 and TWI-6 due to late season green-up. In contrast, the cover of milkweed dropped sharply in these two wet regions. Orchard and grape experienced little to moderate variation across the growing season.

**Fig. 12.**
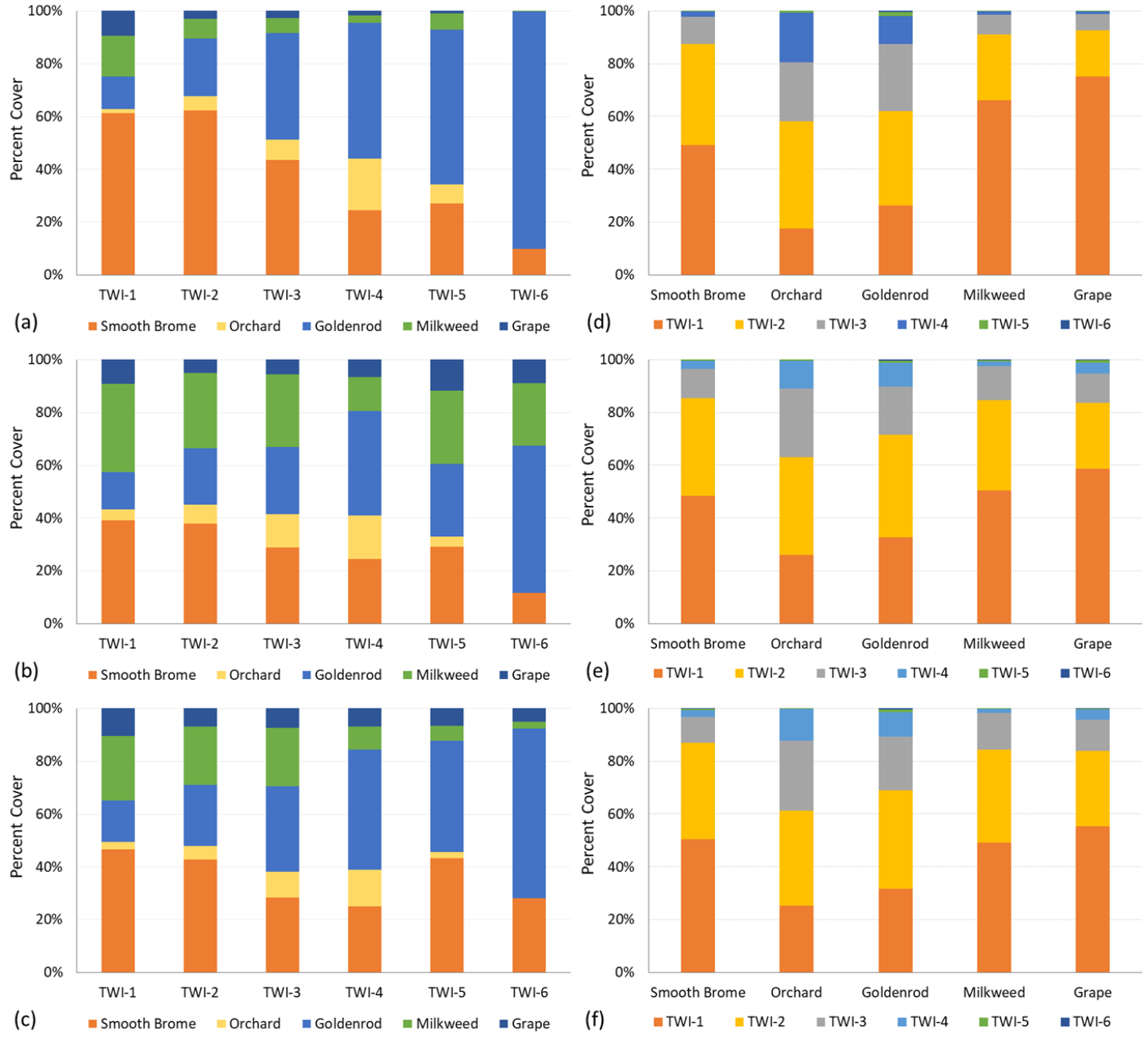
Species percentages in different TWI categories. (a), (b), (c) are species percent cover over the total area of all species in each wetness category in June, August, and September, respectively. (d), (e), (f) are species percent cover in each category over its total area in the entire site in June, August, and September, respectively.

The composition and distribution of native and invasive species across TPI and TWI categories are demonstrated in Fig. 13. Invasive species were dominant across all topographic regions in June, followed by a decrease in August and an increase in September. Despite the seasonal variation, invasive species remained dominant in ridges as well as a large proportion of the site’s mid-slope and upper-slope regions. Considering soil water availability, invasive species dominated drier areas across the growing season, while native species were more likely to reside in wetter regions.

**Fig. 13.**
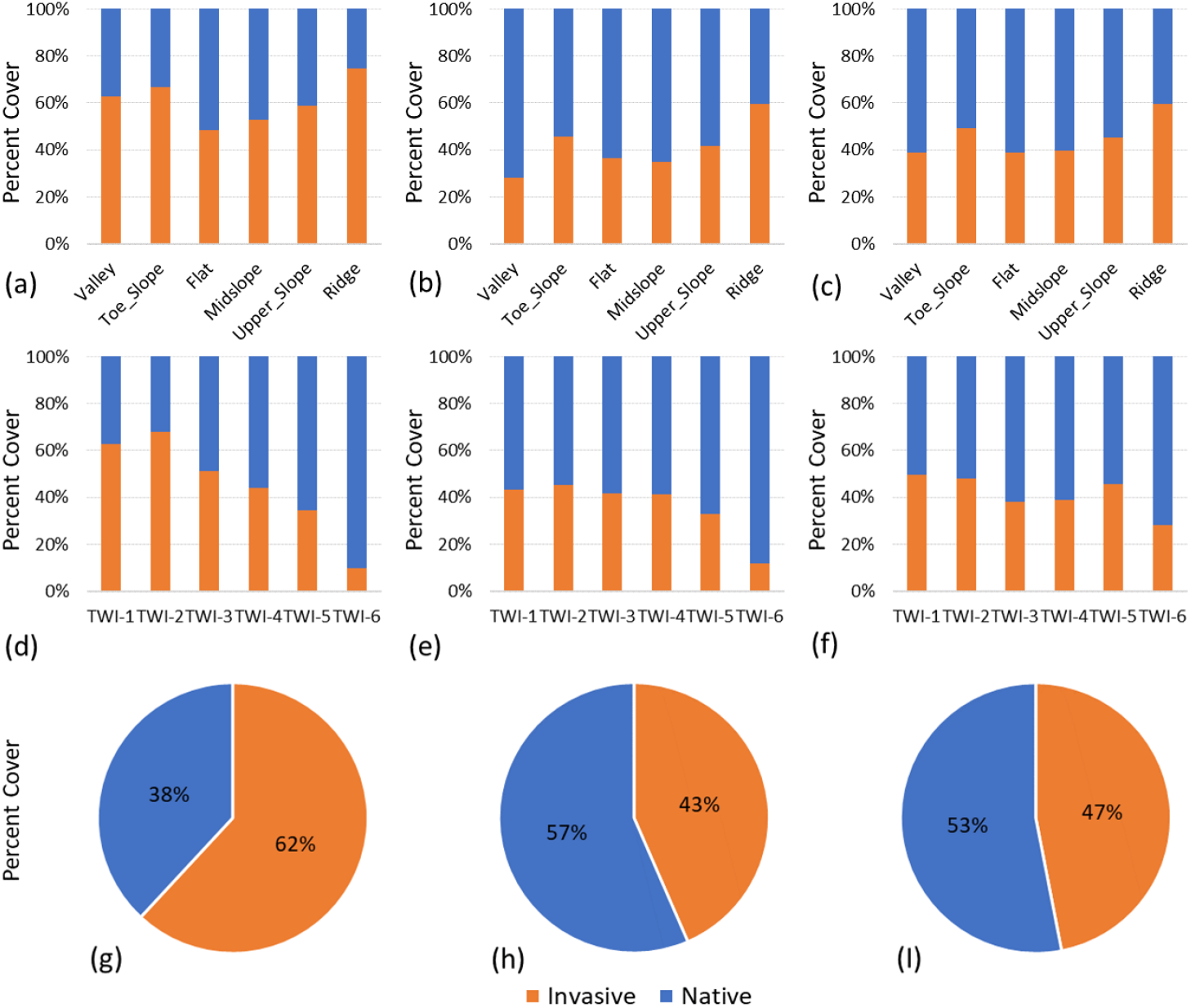
Composition of native and invasive species over different TPI (top figures), TWI (middle figures), native/invasive species cover (bottom figures) in June (left), August (middle), and September(right) in the entire study site.

Overall, the distribution, composition, and variation of the native and invasive species were not only controlled by the seasonality but were also affected by the combined effects of topography and soil water availability. The invasive smooth brome grew early in the season and dominated almost every region in the study area. At the peak growing season, native species (i.e., grape, goldenrod, and milkweed) seemed to prefer wet, lowland, and flat regions, while smooth brome outcompeted native species in upland, steep slope, and dry regions. This invasive species tend to be more tolerant to drought and tend to occupy disturbed areas to acquire resources that are less favourable to native species (Saeidnia et al., 2017). In the late growing season, once native species were senesced, smooth brome re-occupied the flat and wet regions, presumably from a late-season green-up.

Alarmingly, despite seasonal, topographical, and hydrological variation, invasive species occupied a large proportion of the study site with 62%, 43%, and 47% in June, August, and September, respectively. The dominance and expansion of invasive species are threatening to native species, especially in upland, steep-slope, and dry regions. The overwhelming percentage of invasive plant cover of invasive species in the early growing season would acquire a great amount of nutrients, leading to a nutrient shortage for native species that grow later in the season, especially for grape and milkweed. The grim invasion not only suppresses the growth of native species but also impacts the biodiversity and habitats of associated organisms and species. For example, milkweed is known as a source of nectar for many butterflies, bees, and other insects and pollinators, and is essential for the life cycle of the monarch butterfly as it provides food for larvae (Pocius et al., 2017). The decrease in milkweed may seriously impact the population and distribution of these species and others alike. Hence, mapping, monitoring, and understanding the distribution, timing, expansion, and invasion mechanism of these exotic plants are essential for invasion control and management, and for native ecosystem protection.

## 4 Conclusions

In this study, we combined multi-temporal airborne high resolution hyperspectral remote sensing data, topographic position index, and topographic wetness index to map and quantify the small-scale combined effects of seasonality and topography on the distribution, composition, and dynamics of native and invasive species in a heterogeneous grassland site in southern Ontario, Canada. This research work leads to the following conclusions:

1. There were spectral and textural differences between native and invasive species in association with plant properties. These differences showed the potential of airborne hyperspectral data for native and invasive species classification.
2. Random Forest algorithm with narrow-band hyperspectral data inputs offered higher classification accuracy compared to broad-band multispectral data in this heterogeneous environment with a high degree of similarity in plant species.
3. The distribution, expansion, and competition of invasive species in the area were controlled by topography; however, the control varied at different stages across the growing season.
4. The large proportions of invasive species across the growing season indicated that the invasion was a serious threat to native species and wildlife habitats in the area.

The study sets a methodological basis for detecting, mapping, and analyzing invasive species distribution, expansion, and invasion mechanism using high resolution airborne hyperspectral and topographical data. The study also provides essential information about invasive species distribution, behaviour, variation trend for invasive species control and management to protect native ecosystems and wildlife habitats. However, future study is required to investigate the plant traits and biological and ecological characteristics that drive such vegetation patterns to better understand invasion mechanisms.

## Author’s contributions

P.D. and Y.H. discussed and conceived the idea; P.D. conducted the literature review, collected and analyzed data, and led the manuscript writing with revision from Y.H. and A.A; A.A. helped with part of RTK GPS ground control point collection and the calculation of TPI, TWI, and percent cover.

## Data accessibility

The data will be shared from Dataverse, a data portal at the University of Toronto.

## Acknowledgments

This project was financially funded by the Natural Sciences and Engineering Research Council of Canada (NSERC) through the NSERC Discovery Program (RGPIN-386183 awarded to Dr. Yuhong He), and the Graduate Expansion Fund and Graduate Student Research Awards (awarded to Phuong D. Dao) from the University of Toronto. We also thank Bing Lu for providing some reference ground control points for image classification and validation.

## Appendix A

### Additional figures

**Fig. A1.**
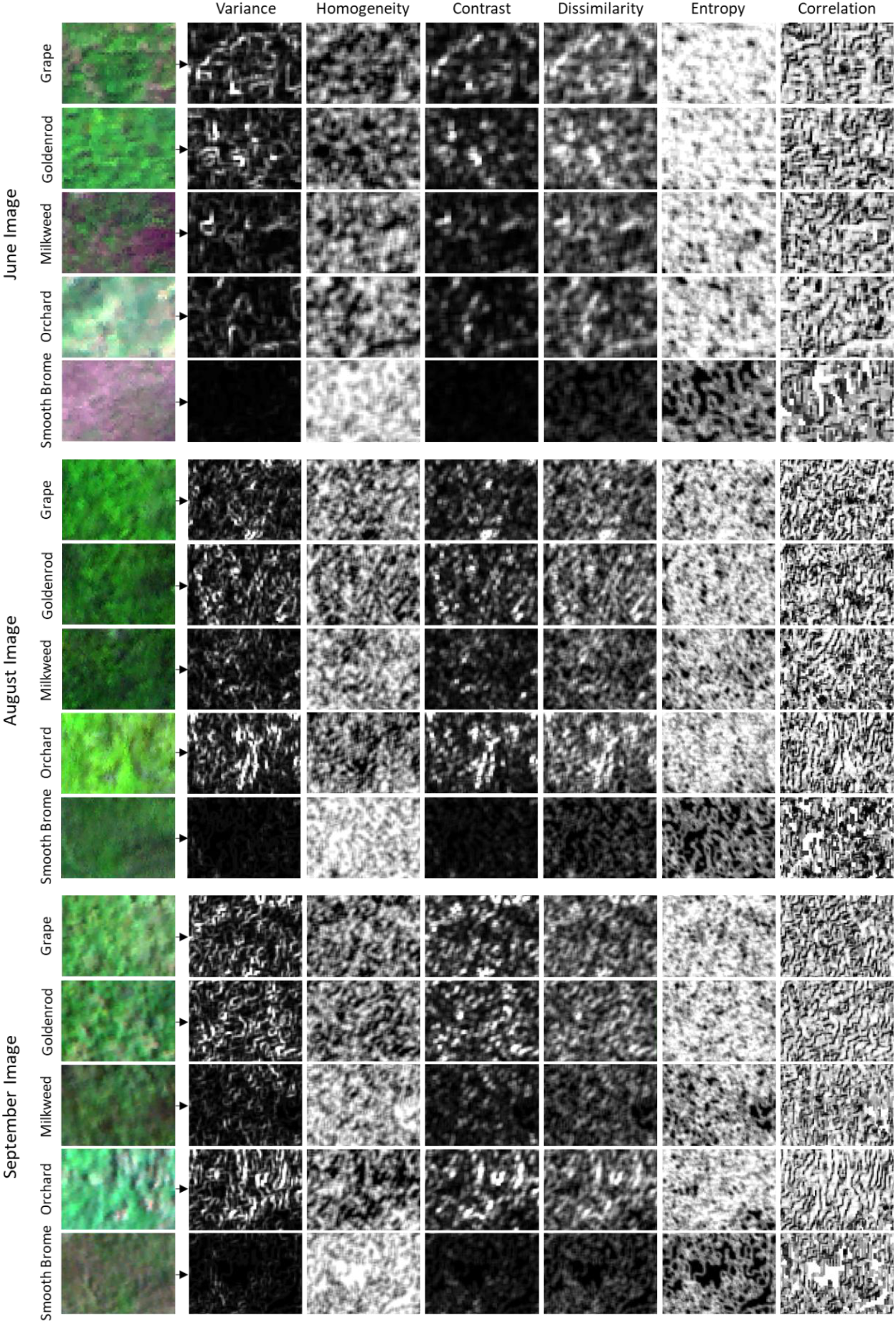
Six textural images of the five species in the study site in June, August, and September.

